# Optimal Control of Spiking Neural Networks

**DOI:** 10.1101/2024.10.02.616330

**Authors:** Tiago Costa, Juan R. Castiñeiras de Saa, Alfonso Renart

**Author notes:** Equal contribution (Tiago Costa, Juan R. Castiñeiras de Saa).

## Abstract

Control theory provides a natural language to describe multi-areal interactions and flexible cognitive tasks such as covert attention or brain-machine interface (BMI) experiments, which require finding adequate inputs to a local circuit in order to steer its dynamics in a context-dependent manner. In optimal control, the target dynamics should maximize a notion of long-term value along trajectories, possibly subject to control costs. Because this problem is, in general, not tractable, current approaches to the control of networks mostly consider simplified settings (e.g., variations of the Linear-Quadratic Regulator). Here, we present a mathematical framework for optimal control of recurrent networks of stochastic spiking neurons with low-rank connectivity. An essential ingredient is a control-cost that penalizes deviations from the default dynamics of the network (specified by its recurrent connections), which motivates the controller to use the default dynamics as much as possible. We derive a Bellman Equation that specifies a Value function over the low-dimensional network state (LDS), and a corresponding optimal control input. The optimal control law takes the form of a feedback controller that provides external excitatory (inhibitory) synaptic input to neurons in the recurrent network if their spiking activity tends to move the LDS towards regions of higher (lower) Value. We use our theory to study the problem of steering the state of the network towards particular terminal regions which can lie either in or out of regions in the LDS with slow dynamics, in analogy to standard BMI experiments. Our results provide the foundation of a novel approach with broad applicability that unifies bottom-up and top-down perspectives on neural computation.

## 1 Introduction

Normative or top-down approaches to neural computation work through the specification of particular cost functions that the dynamics of the network should optimize, and which formalize specific computational goals. The standard approach is to shape the network dynamics by finding appropriate values for the weights of a recurrent neural network (RNN) of firing rate units [1, 2], although progress has also been made towards shaping the dynamics of networks of spiking neurons [3, 4]. This research program has been successful at revealing how RNNs can be made to embody a series of dynamical motifs that constitute specific implementations of standard computations needed to solve a host of behavioral tasks studied in the laboratory [5, 6].

Despite its success and popularity, however, there are shortcomings to this approach. Functional constraints shaping the structure of these networks do not appear to be sufficient to drive the emergence of large-scale anatomical structure. Trained networks don’t typically display modularity or develop spatially confined functional specializations – unlike brains – although some studies are starting to explore modular structures [7, 8]. In contrast, brain architecture is modular and hierarchical, with high-level executive areas shaping activity of hierarchically lower areas [9, 10]. Such top-down projections enforce that processing is adapted to the current context [11, 12, 13] – which varies on fast time-scales – whereas the local connectivity of lower level areas is often understood to reflect the statistical structure of the environment, in the sensory domain [14, 15], or the implementation of motor-primitives or hard-wired action policies in the motor domain [13, 16] – which are stable over much longer time-scales.

Architectures of this type have been suggested to implement cognitive control [17, 18, 19, 20]. Our working hypothesis is that an essential feature of such architectures is that rapid changes in functionality needed to drive context-appropriate behavior are too quick to rely on learning and thus reflect changes in the dynamics of a modular system. Although it is currently not understood how these modular control architectures work, a pre-requisite would be to understand the optimal external input that a controller should use in order to control the activity of a neural network – whose default dynamics are defined by a given, in principle task-independent, recurrent connectivity – so as to meet a specific control target. Here, we develop a framework for solving this problem within the context of optimal control theory [21, 22]. We consider networks of stochastic spiking neurons [23, 24] with low-rank connectivity. Low-rank connectivity induces low-dimensional dynamics which are computationally interpretable [25, 26, 27], and provides a good test bed for studying how external inputs may modify the locally induced dynamics of a network when appropriate.

Formalizing this problem rquires defining a notion of value over instantaneous and terminal network states and a cost of control that penalizes large deviations from the default dynamics of the unperturbed network. The goal of the optimal control is to steer the dynamics of the network to maximize expected future value. The most thoroughly studied situation corresponds to the so-called linear-quadratic regulator (LQR), in which the dynamics are assumed to be linear and the instantaneous costs are quadratic [21, 22]. In this case, the optimal control law is analytically tractable, but the computational power of linear networks is limited. Variants of this problem have been used to study control of neural activity in neuroscience [28, 29, 30]. A number of methods exist to consider non-linear dynamics [31, 32, 33]. Deep reinforcement learning (RL) has also been used for continuous control [34], showing that accurate control is possible for large dimensional state spaces. Here, we formalize the problem of optimally controlling the activity of a network of spiking neurons with low-rank connectivity, and solve it to study the phenomenology of the control problem for a case study of a rank 1 network.

## 2 Problem formulation

We consider a network of *N* neurons whose membrane potential evolves according to

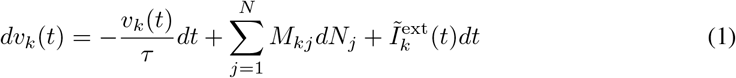

where *dN*_*j*_ is the spike train of neuron *j, M*_*kj*_ are the elements of the recurrent connectivity, and 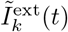 is an external input. Neurons fire spikes as inhomogeneous Poisson processes with an intensity function 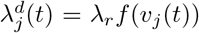 where *f* (.) is an invertible non-linearity and λ_*r*_ is a gain factor with units of Hz. The previous equation can be integrated in time to obtain **v**(*t*) = **Ms**(*t*) + **I**^ext^(*t*), where 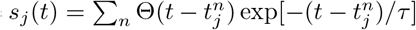 and 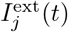 are the low-pass filtered spike trains and external input respectively (i.e., 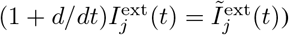. We generally consider static external inputs, since time-varying inputs will be provided by the control. The effective stochasticity of this network can be controlled by the shape of the non-linearity [23]. In the deterministic limit, one recovers a network of leaky integrate and fire neurons [35, 3].

The connectivity matrix 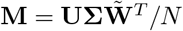 is of rank *K* ≪ *N* (Fig. 1a), where **U** and 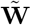 are *N* -by-*K* matrices with *O*(1) components and orthogonal columns, i.e., 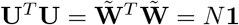. We write 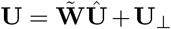 where 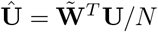 and **U**_⊥_ is the projection of **U** onto the subspace orthogonal to 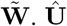 measures the alignment between the row and column subspaces of the connectivity, which determines the qualitative nature of the recurrent transformation [25, 26]. In our case, it determines how much of the network’s output is fed back into the input subspace to which the network is sensitive to. We will work in an orthogonal basis given by the columns of 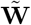 and additional orthogonal vectors as needed to span **U**_⊥_, which we arrange as the columns of a matrix **W**_⊥_, i.e., 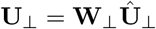, with 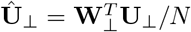. The total dimension of this basis set is between *K* and 2*K*, depending on the alignment between **U** and 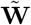, but we take it to be 2*K* for our derivations. The low-dimensional dynamics of the network is characterized by the value of the overlaps between the filtered spike trains and 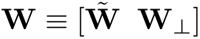^1^

**Figure 1:**
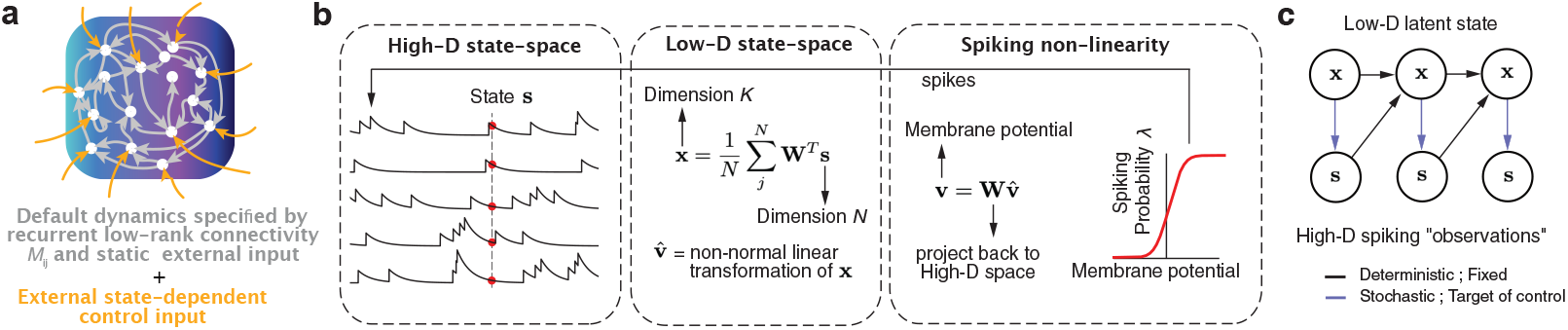
Recurrent spiking network. **(a)** Schematic description of the control problem. **(b)** Schematic description of spiking recurrent network dynamics. The high-dimensional membrane potentials which determine the spiking activity are a linear transformation of a low dimensional latent variable 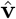, whose value depends on the overlaps **x** between the filtered spike-trains **s** and a series of vectors that span the low-rank connectivity matrix. **(c)** Generative model.

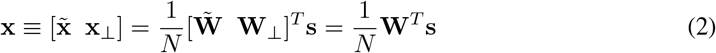

Denoting by **s**_⊥_ the part of **s** orthogonal to **x**, and assuming that the external input projects into the low-dimensional subspace, i.e., 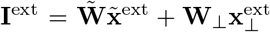, we can write the equation for the membrane potentials as

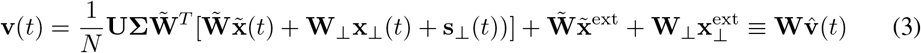

where 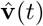 is the time-varying low-dimensional latent variable which completely determines the membrane potential of the neurons of the network (Fig. 1b), given by

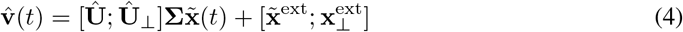

### 2.1 The control problem

In our formulation, (stochastic) action policies correspond to spiking probabilities. As mentioned above, the default policy corresponding to the uncontrolled network dynamics is given by

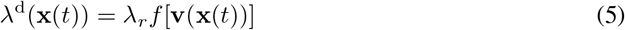

Thus, at each time *t*, each neuron has two available “actions”: to spike or to remain silent, which have probabilities λ^d^*dt* and 1 − λ^d^*dt* under the default policy. Since the (high-dimensional) spiking probability depends exclusively of the (low-dimensional) overlaps – through the membrane potentials in Eq. (3) – the dynamics of the overlaps completely characterize the temporal evolution of the network’s state

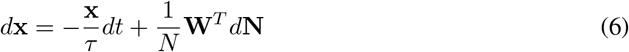

Equations (5) and (6) contain the same information as the policy *π*(*a* | *s*) and state-transition matrix *T* (*s*′| *a, s*) in a standard tabular RL setting. As is standard, the evolution of the state given the current state and action (i.e., the temporal evolution of the low-dimensional network state given the spikes – which is in fact deterministic in our framework) is not controlled (Fig. 1c). Instead, we want to modify the policy, i.e., the spiking probability given **x**(*t*), in order to maximize the following expectation

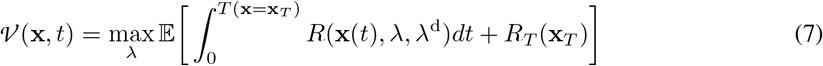

The quantity *𝒱* (**x**) is the value function of the problem. Given an instantaneous ‘reward rate’ *R*(**x**(*t*), λ, λ^d^) and a ‘terminal reward’ *R*_*T*_ (**x**_*T*_) obtained when reaching a terminal region **x**_*T*_, the value function is equal to the expected future reward acting under the optimal policy. Since we want the control to modify the spiking probability of each neuron to leading order, we consider a finite instantaneous reward rate per neuron

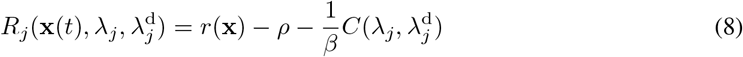

and thus the total instantaneous reward rate is extensive^2^: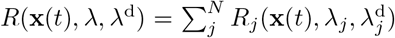. The quantities *R*_*T*_ (**x**_*T*_) and *r*(**x**) are state reward/costs, i.e., value that is experienced by virtue of the network being at some particular state. For instance, in the context of a BMI experiment, the goal might be to reach a particular magnitude for a readout projection, which would determine **x**_*T*_, and *R*_*T*_ (**x**_*T*_) would be the reward for reaching that location. *r*(**x**) can be used to represent sensory feedback on the way to the goal, and *ρ* is a cost of time. 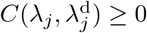 represents the cost of control, i.e., a penalty that is paid by using policy λ when the default policy is λ^d^, and thus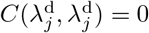. Its form will be discussed below. Finally, the parameter *β* measures the sensitivity of the agent to the cost of control. When *β* ≫ 1 control is cheap, and the optimal policy will take the network’s state directly and quickly to the target **x**_*T*_. When *β* ≪ 1, control is costly, and the control policy will try to deviate from the default dynamics as little as possible. This situation is the most interesting, as it allows exploring how to best use the default dynamics to accomplish a given task. We will refer to *β* as the ‘control gain’.

To proceed, we use Kolmogorov’s backward equation [36, 37] to obtain a differential representation of the problem^3^

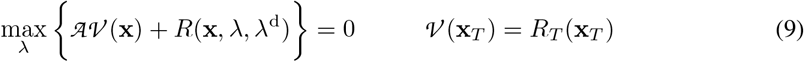

in which *𝒜* is the infinitesimal generator of the process **x**(*t*). For our jump process, we obtain the following expression, which is the Bellman Equation for our problem

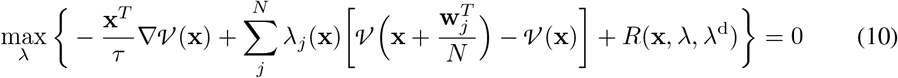

where 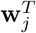 is the *j*^*th*^ column of **W**^*T*^ in Eq. (6).

### 2.2 Conjugate control

In order to proceed, one needs to specify the control cost 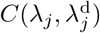. We initially consider 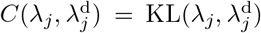, the Kullback-Leibler divergence between the two policies, as in the framework of KL control [38, 39, 40]. It is straightforward to show (Supplementary Materials) that in this case, the optimal policy is given by

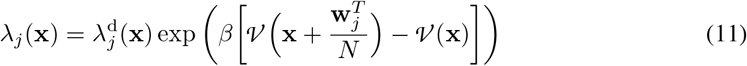

Control, therefore, modifies the policy through a state-dependent gain factor. However, if the spiking non-linearity *f* (.) is exponential, then we can write

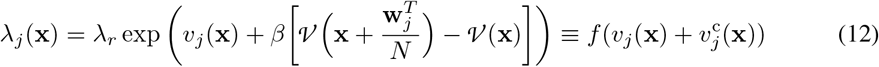

When written in this form, the optimal control law can be interpreted as as additive synaptic input 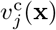 which could presumably come from another network playing the role of a ‘controller’. Furthermore, the optimal control input is interpretable. Recall that 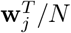 measures how much the network state **x** changes upon a spike from neuron *j*. Thus, the optimal control policy excites (inhibits) a neuron to the extent that a spike from that neuron (relative to no spiking) moves the network state towards a region of higher (lower) value on the low-dimensional manifold. This difference in values is equal to the *advantage* of the spike [41].

An exponential non-linearity, however, is somewhat unsatisfactory, since it’s unbounded and numerically unstable. We realized, however, that it is possible to retain an interpretable optimal control law consisting of an additive external synaptic input equal to the spike’s advantage, using any invertible non-linearity, as long as the functional form of the control cost is selected appropriately. In particular, we *impose*

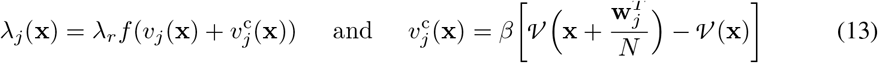

but let *f* (.) be any invertible non-linearity. To find the maximum in Equation (10), we take derivatives^4^ with respect to λ_*j*_, obtaining

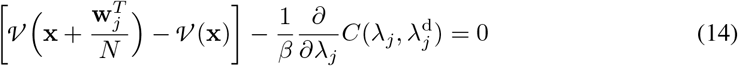

From Equations (13) and (14) we thus obtain

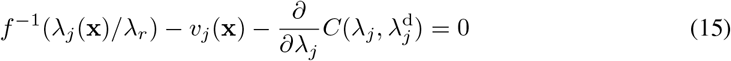

This is an ODE in the unknown function 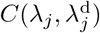 which can be solved with the boundary condition 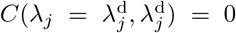. We solved this equation for a sigmoidal non-linearity 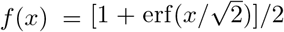 (Supplementary Materials). The resulting costs are convex and qualitatively similar to a KL divergence. Since the functional form of the non-linearities in the default and optimal policies are equal in this framework, we refer to this as conjugate control, in analogy to Bayesian inference.

To find a more compact form for the Bellman Equation, we Taylor-expand the advantage, obtaining 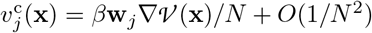. Thus, to leading order, the control input is given by **v**^c^(**x**) = **W***β*∇ *𝒱* (**x**)/*N*, showing that the low-dimensional control on the manifold is proportional to the gradient of the value. Grouping terms in Equation (10) one can then write the following general form

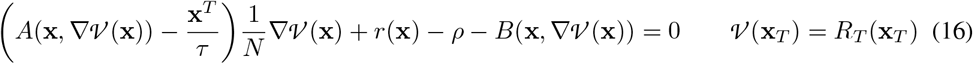

Where

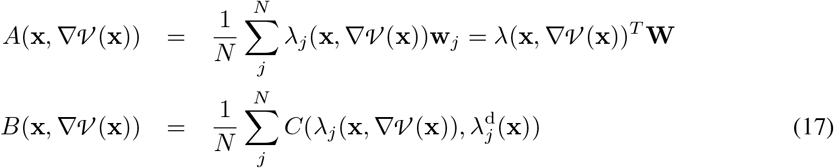

*A*(**x**, ∇ *𝒱* (**x**)) is a column vector of size 2*K* with elements equal to the overlap between the firing probabilities and each column of **W**, and *B*(**x**, ∇ *𝒱* (**x**)) is equal to the average control cost at **x**. Note that all quantities in Equation (16) are intensive, and the value function only appears through *𝒱* (**x**)*/N*. Thus, if *R*_*T*_ (**x**) ∼ *O*(*N*), then the value function will be extensive. Although *𝒱* (**x**) ∼ *O*(*N*), because each spike only leads to a change *O*(1*/N*) on the collective macroscopic state **x**, the advantage of each spike is ∼ *O*(1) and thus the control input and the default input from the recurrent network have the same asymptotic scaling, making the control problem non-trivial. To focus on this asymptotic regime, we redefine *R*_*T*_ (**x**_*T*_) → *NR*_*T*_ (**x**_*T*_) and thus drop all factors of *N* from the Bellman equation and boundary conditions.

For an arbitrary low-rank connectivity, the sums in Equations (17) have to be computed explicitly. However, if the elements of **U** and 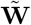 are assumed to be Gaussian, then for both the exponential and sigmoidal non-linearities we consider, the sums can be computed analytically for large *N* as expectations over the distribution of the weights, which renders the Bellman Equation truly low-dimensional. Details of the derivation for both non-linearities can be found in the Supplementary materials.

## 3 Case study for a rank-1 network

We applied our theory to study the phenomenology of control for a rank 1 network of neurons with a sigmoidal spiking non-linearity. The control policy is completely specified by the value function (Equation (13) (asymptotically it is proportional to its gradient), and the value is the solution of the Bellman Equation (Supplementary Materials, Equation (40)). We used an adaptation [42] of the Deep Galerkin Method [43] to solve this non-linear PDE (Supplementary Materials). In Figure 2 we consider a rank 1 network with 2 nearby stable fixed points separated by a saddle, in such a way that the dynamics between the fixed points is slow (Figure 2a; parameters for all figures can be found in 1 in the Supplementary Materials). Because the network is rank 1, the full dynamics is two-dimensional.

**Figure 2:**
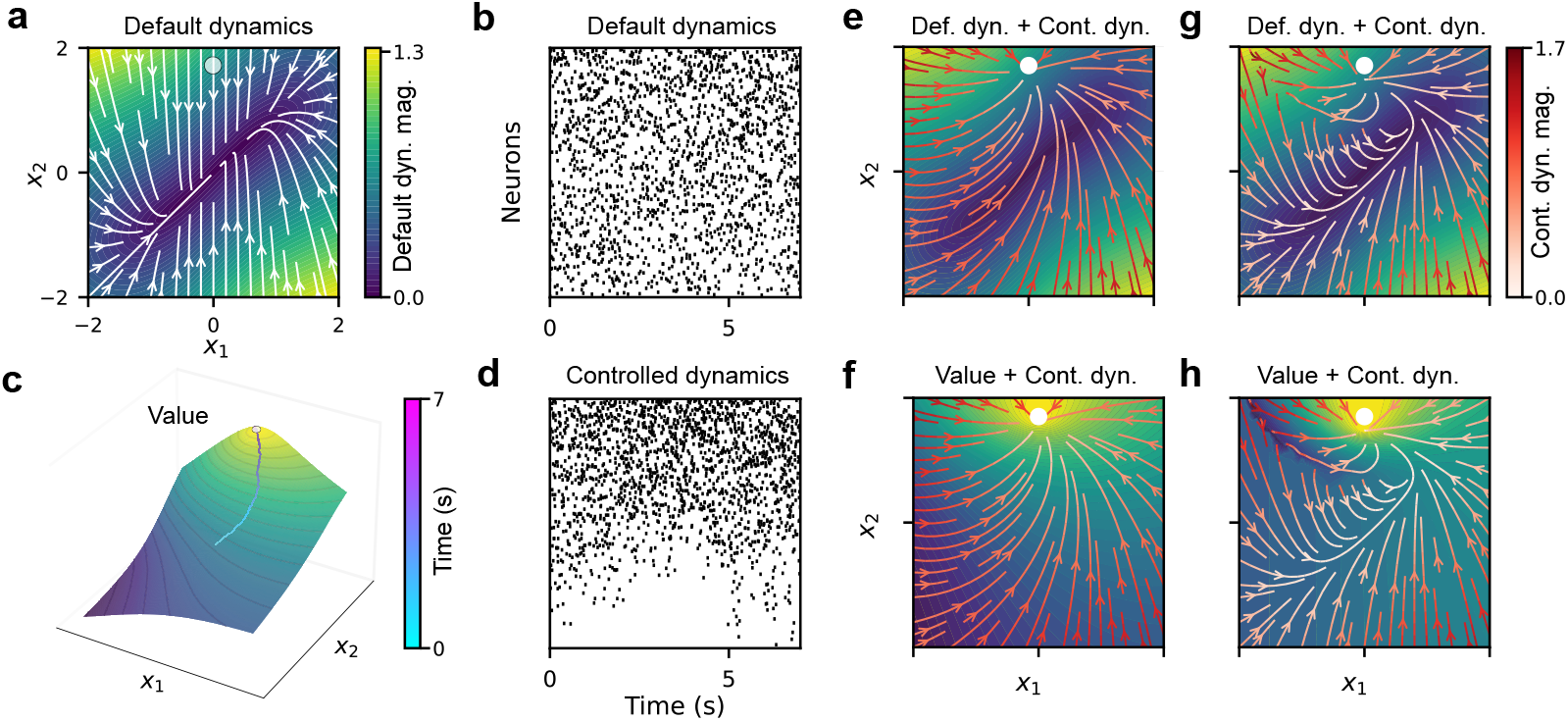
Controlling a rank 1 network. Effect of control gain. **(a)** Default dynamics considered for this example, displaying a slow manifold aligned to the diagonal. The control problem in **b-h** is to take the network state to the circle in the top-middle of the panel. **(b)** Example raster of a simulated network with 500 neurons following these default dynamics. **(c)** Value function for high *β* corresponding to reaching a small circular region, with an example controlled trajectory shown on top of the surface. The control input is proportional to the gradient of the Value at each point of the trajectory. The color represents the time along the path, with the target being reached at approximately 5*s*. **(d)** As in **b**, but corresponding to the example controlled trajectory in **c. (e, f)** Controlled dynamics for the setting in **c**, with the streamline color representing its magnitude. On top, the background represents the magnitude of the default dynamics as in **a**, and on the bottom it shows the Value function. **(g, h)** Same as **e, f**. but for a small value of *β*.

The control problem is to drive the newtork state **x** to a circular target (dotted circle in Figure 2a). We assume that the only non-zero reward/cost is the cost of time, so the control wants to take the network to the target as soon as possible. The value function thus approaches zero at the target and decays away from it (Figure 2b). Although the value function is solved in the asymptotic limit, we can use it for a finite network. We considered a network of 500 spiking neurons (spike raster for default dynamics in Figure 2c). As expected, starting from **x**(*t* = 0) = [0, 0], the control took the network to the target (Figure 2d,b). In Figure 2e-h we explore the role of the control gain *β*. When *β* is larger (same value as in Figure 2b), the controlled trajectories go relatively directly to the target (Figure 2e,f). Thus, for large control gains, the control overcomes, and is relatively insensitive to, the default dynamics. When *β* is smaller, the dynamics are more interesting, as the control policy tries to use the default dynamics as much as possible to take the state to the target. The optimal strategy is to allow the state to reach the slow manifold using the default dynanics, bring it close to the target there (paying little control cost), and only spendig a significant control cost when close enough to the target (Figure 2g,h). For low control gains, the dynamics is much richer. Notice for instance the appearance of a separatrix (i.e., a ‘decision bound’) by the lower right corner of the state space, which partitions the regime where it pays off to drive the state directly to the target (low cost of time, high cost of control), from the regime where it pays off to let the default dynamics do most of the work (high cost of time, low cost of control).

In order to highlight that the optimal controlled dynamics, for the same control targets, depends strongly on the default dynamics if the control gain is low, we studied the same control problem, for two different rank 1 networks. The first one has a single stable fixed point and roughly isotropic dynamics around this fixed point (Figure 3a). The second one is the same one studied in Figure 2. As typical in brain-machine interface (BMI) experiments, we derive optimal policies for 8 targets lying in a circle in the two-dimensional state space. In the first case, all the control problems are roughly symmetric with respect to the fixed point structure of the default dynamics, the controlled trajectories are straight to the targets, and the value of the initial state is also target-independent. In the second case, the default dynamics strongly shape the optimal policy, so the trajectories to the different targets show different amounts of curvature, and the value of the initial condition (which partly shows how long it is going to take to reach the target) also varies strongly for different targets. In BMI experiments involving controlling the position of a cursor on a screen, it is often observed that cursor trajectories are not straight. Our results suggest that the shape of these trajectories is not idiosyncratic, and instead might reflect compromises in the ability of the rest of the brain to control the activity of a local network with its own intrinsic manifold. Steering the activity of a local network away from regions favored by its intrinsic dynamics might be more difficult, or even impossible in some cases [44].

**Figure 3:**
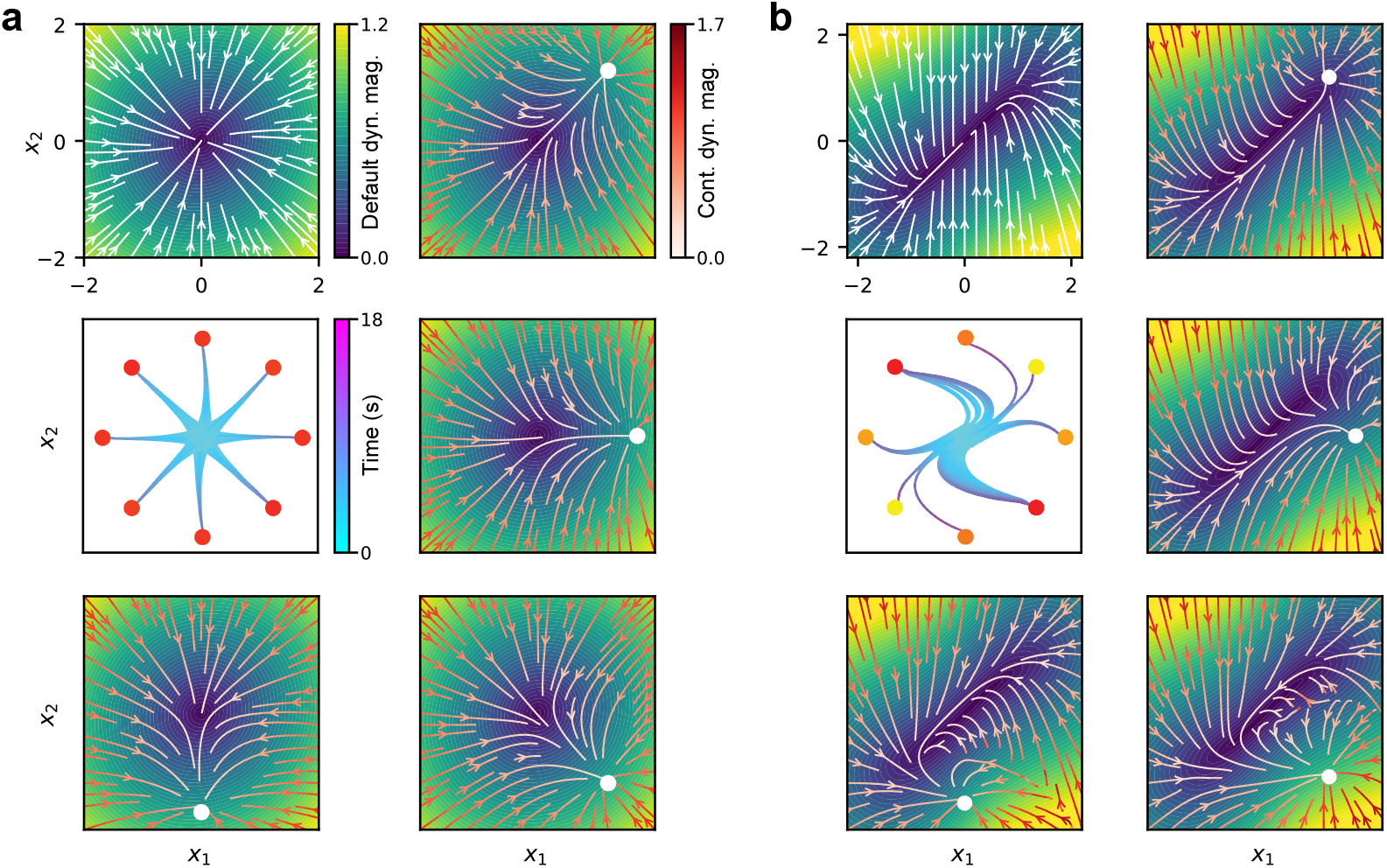
Controlling a rank 1 network. Effect of default dynamics. The network has to reach one of the 8 available targets in the LDS, and we are showing the behavior under two different default policies. **(a)** Case study with an isotropic default policy (up-middle). From up-right to bottom-middle, we show the controlled dynamics (magnitude as streamline color) that reaches to the corresponding target (due to symmetry, the remaining targets just involve a rotation). The background remains the magnitude of the default dynamics. The center shows example trajectories reaching to each target, spanned from a circle around the origin. The color of the targets indicates their value, and the color of the lines means time along the trajectories. **(b)** Same as **a**, but for a default policy that features a slow manifold aligned to the diagonal (same as in Figure 2). All colormaps are common.

## 4 Limitations and future directions

Naturally, given that we have only just established the feasibility of our framework, a number of important issues remain to be addressed. One limitation is the absence of an unstructured part of the connectivity, as is common in network models [45, 25]. If the variance of the unstructured part of the connectivity is large enough, time-dependent chaotic activity develops [45, 24]. In our framework, when the elements of the low-rank connectivity matrix are ∼ *O*(1*/N*), the dynamics is assymptotically deterministic. Random connectivity of sufficient variance would lead to diffusive dynamics. Although in principle, this does not represent a fundamental challenge for control, the exact consequences of this stochasticity in the trajectories remains to be evaluated. Another biophysical constraint we have not yet considered is separate excitatory and inhibitory neurons, which have strong implications for the dynamics of recurrent circuits [46, 47, 48, 49].

We have started by considering first-exit problems, but some important control problems are not naturally construed as first exit. Some examples include control targets corresponding to particular dynamical regimes [50], or control targets that correspond to the maximization of variance in neural along particular dimensions, which might be important for controlling the flow of information across brain areas [20].

Another area of interest is time-dependent problems. These arise naturally in the study of inference and decision-making. Indeed, the value function for solving a classical two-alternative foced-choice decision problem depends explicitly both on accumulated evidence and on elapsed time [51, 52]. We are particularly interested in control policies that implement efficient decision-commitment strategies [52], and also in the problem of whether controlling the activity of a network so that it can optimally categorize stochastic inputs, will automatically lead to network dynamics that effectively compute posterior beliefs.

Finally, we would eventually like to explictly model the controller as a spiking neural network itself. The framework of spike coding networks [3, 29] seems like an interesting avenue to consider, since these newtorks can implement non-linear functions [4] such as our state-dependent policies. An actual implementation of the full plant-controller system would also allow investigating important questions such as imperfect state-estimation or the presence of delays in the feedback loop.

## 5 Discussion

We have developed a mathematical framework for optimal control of stochastic spiking low-rank recurrent neural networks. Our framework differs from standard implementations in that we don’t assume a particular parametric form for the controller. Instead, we formally optimize the stochastic transformation from membrane potentials to spikes. In addition to describing spiking activity, this choice results in principle in a more general (less constrained) optimal control law. However, in general, this will lead to solutions that are more difficult to interpret biologically, as control is naturally construed to be an additive external synaptic input from a different brain area which is then acted upon by a control-independent spiking non-linearity. However, inspired by the special role of an exponential non-linearity in KL control, we developed the concept of conjugate control, which allows the control input to be aditive under a fixed non-linearity, while still allowing for un-parametrized control policies.

The resulting optimal control is nicely interpretable as being proportional to the advantage [41] of a spike, i.e., an external input that excites or inhibit neurons depending on whether their spiking moves the low-dimensional network state twowards regions of higher or lower value.

To obtain the value function, we derived a Bellman Equation for the problem – taking the form of a challenging non-linear PDE, which we solved using a deep neural network [43, 42]. Although this method works and can scale up to approximately 20 dimensions [42], it is not clear how such policies could be learned. Although learning remains an open problem, we have already started to explore alternative ways to derive the value function. In particular, we developed an on-policy iteration method that computes the value by evaluating the accumulated cost along simulated trajectories of the dynamics (see Supplementary Materials for a proof of principle of this method), analogous to the path-integral method for control [38, 39]. A method of this sort using ‘roll-outs’ [53, 54] seems much easier to implement by gradual learning in neural tissue. Additionally, it would be interesting to explore methods that find the optimal policy directly without going through the value function [55]. Although the fact that the optimal policy is the gradient of the value is intuitively reasonable, calculating this gradient is an extra computational cost, and requires extending the domain over which the value is calculated. Ideally, in high-dimensional problems, one would like to only calculate the optimal policy for the small-region of states that are frequently visited by the network.

We have applied our theory to the study of first-exit problems that resemble those in BMI experiments, where the activity of a local network needs to be steered towards a target. Several studies have modelled BMI control recently [56, 57], with an emphasis on describing limitations in learning derived from the relationship between control targets and the network’s intrinsic manifold [44] – an issue we addressed indirectly in our results. However, these studies did not emphasize feedback control [57] or even dynamics [56]. In contrast, our goal is to characterize in detail how to optimally dynamically control the activity of recurrent spiking networks. Our results show that, when the control gain is low, the control policy interacts with the default dynamics in interesting ways, giving rise to bifurcations that reflect trade-offs between the different costs that are part of the optimization problem.

It is fair to ask whether studying optimal control policies might not be overambitious, given the many mechanistic constraints faced by neural control systems. We believe, however, that precisely by understanding how the structure and dynamics of particular circuits shapes the solution of given computational problems that operate through the control of such circuits, it will become possible to systematically evaluate the impact of the mechanistic contraints that different control architectures impose on function.

## A Supplemental material

### A.1 Optimal policy for KL control

The fact the optimal policy in KL control is given by a multiplicative gain factor – equal to the exponential of the advantage multiplied by the control sensitiviy *β* – is well established, in general for discrete policies [39], and also for policies in continuous time [52]. Here we provide that can be generalized to other functional forms for the control cost.

Consider the KL divergence between two Poisson rates λ(*t*) and λ^d^(*t*) at a particular time

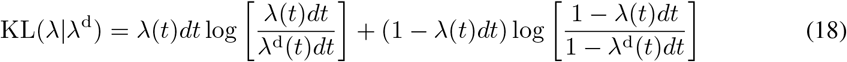

Developing to first order in *dt*, one obtains

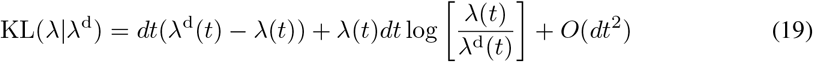

Thus, under KL control, our control cost 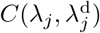 for the spiking of neuron *j* per unit time is given by

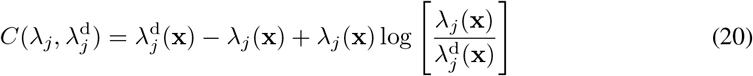

To obtain the optimal policy, we plug in this expression into Equation (10)

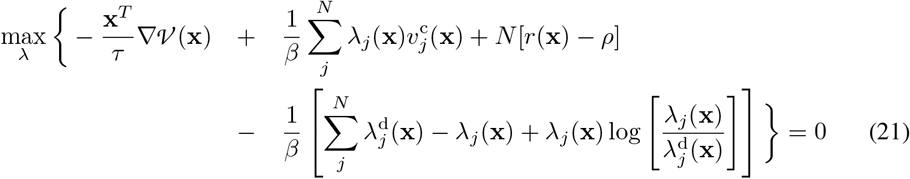

where 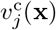 is defined in Equation (13). Taking out the terms of the equation that don’t depend on λ

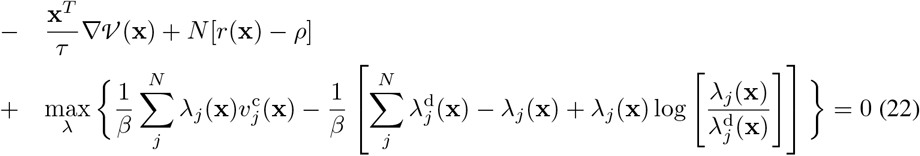

such that

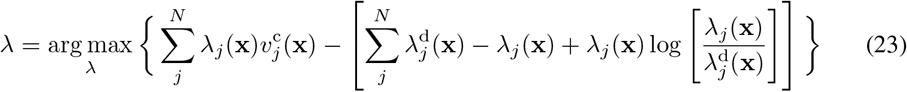

As mentioned in footnote (4), Equation (23) can be solved by simply taking derivatives with respect to λ and setting them to zero. In addition, the equations for each neuron *j* decouple, so we obtain

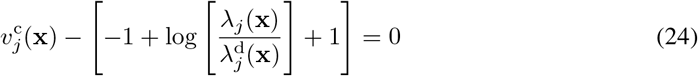

so that

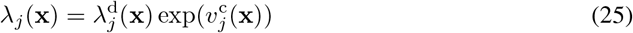

which is Equation (25)

### A.2 Cost of control for sigmoidal non-linearity

Consider 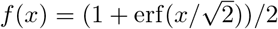. For this non-linearity, Equation (15) reads

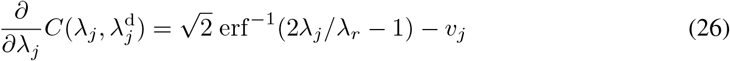

The solution of this equation in terms of *v*_*j*_ and 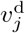 is

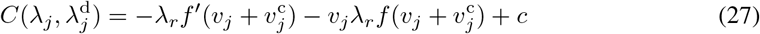

where 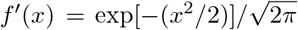 and *c* is an integration constant. Solving for *c* by imposing 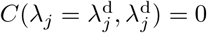, one obtains

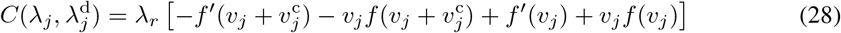

This expression is useful to compute the Gaussian integrals in section A.4, but one can also write the cost in terms of only λ_*j*_ and 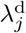

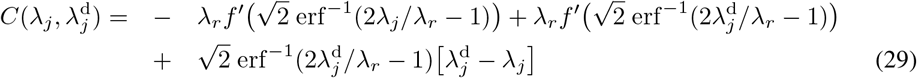

This function is plotted in Figure X

### A.3 Bellman Equation for KL control

Our goal is to evaluate *A*(**x**, ∇ *𝒱* (**x**)) and *B*(**x**, ∇*𝒱* (**x**)) in Equations (17), in order to have a closed form expression for the Bellman Equation (16). In this section 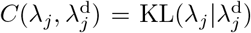 and the spiking non-linearity is exponential, i.e., *f* (*x*) = exp(*x*). We assume the value function is extensive, so we drop all the *N* factors in terms proportional to ∇ *𝒱* (**x**)*/N* in Equation (16). Using Equations (20) and (25), the control cost *B*(**x**, ∇ *𝒱* (**x**)) reads in this case

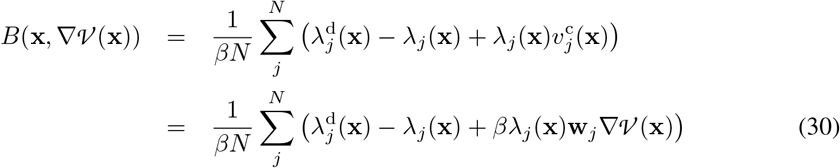

Thus, for the case of KL control, there is a cancellation between the last term in the control cost, and the first term, proportional to *A*(**x**, ∇ *𝒱* (**x**)), in the Bellman Equation (16). After this cancellation, the equation reads

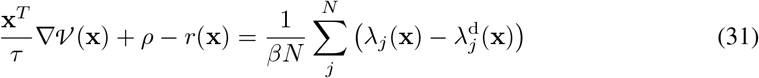

Consider 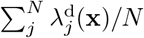, i.e., the average firing probability at **x** in the uncontrolled network. These population averages can be calculated analytically if the elements of the columns of the matrix **W** specifying the low-rank synaptic connectivity are independent Gaussian variables (which, for 13 simplicity, we take to have zero mean). For the exponential non-linearity associated to the KL control cost, one can then evaluate this sum as

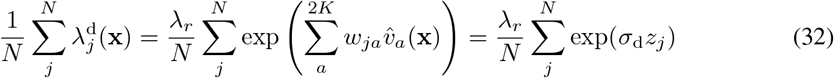

where 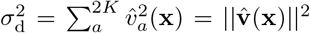 and the *z*_*j*_ are standard normal variables. For large *N* this sampling-based expectation can be well approximated by the expectation over the distribution of *z*_*j*_

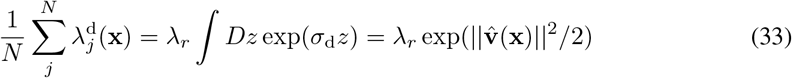

where 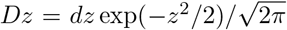 and where the integral is done by completing the square. The calculation of 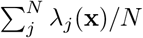 can be performed identically, recalling that

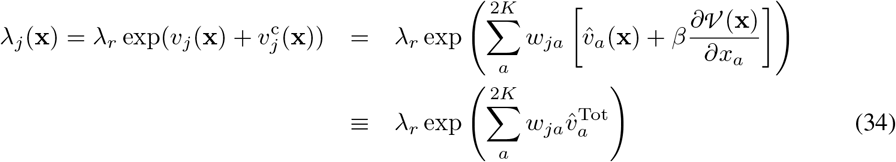

where we have defined 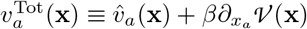, which is equal to the total low dimensional latent state for the network under the controlled policy. Replacing these expressions in Equation (31), one obtains the following expression for the Bellman equation

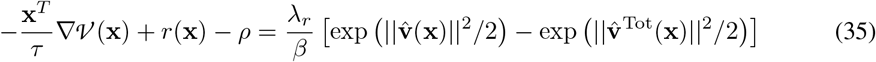

This equation needs to be solved with boundary condition *𝒱* (**x**_*T*_) = *R*_*T*_ (**x**_*T*_).

### A.4 Bellman Equation for sigmoidal non-linearity

In this section, we evaluate *A*(**x**, ∇ *𝒱* (**x**)) and *B*(**x**, ∇*V* (**x**)) assuming that 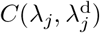 is given by Equation (28), and that 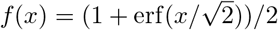 (so that 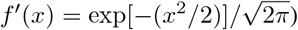. We start by calculating

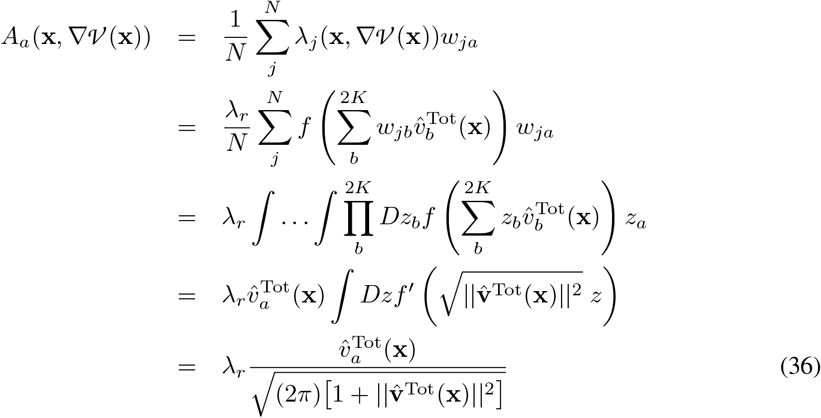

where we have used the identity 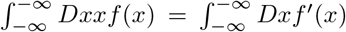 (a particular case of Stein’s lemma) in passing from the 3^*rd*^ to the 4^*th*^ lines, and where the integral in the 4^*th*^ line can be done by completing the square, since *f* ′(*x*) is Gaussian. In sum

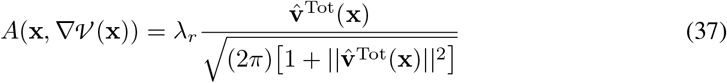

In order to calculate *B*(**x**, *𝒱* (**x**)) we need to evaluate four summations corresponding to each of the four terms in Equation (28). Each of these requires the same identities for Gaussian integrals used in the previous calculations. Thus, we only give the results

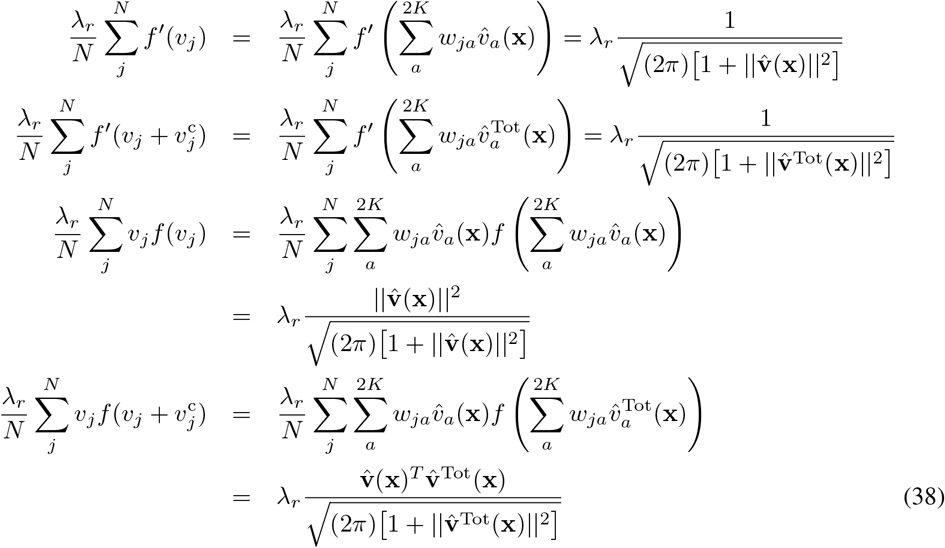

thus, collecting terms

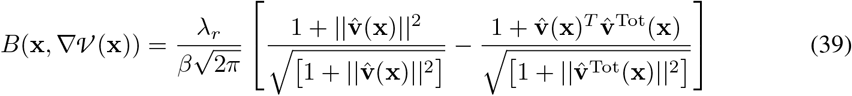

Therefore, the Bellman Equation for the error-function sigmoidal non-linearity reads

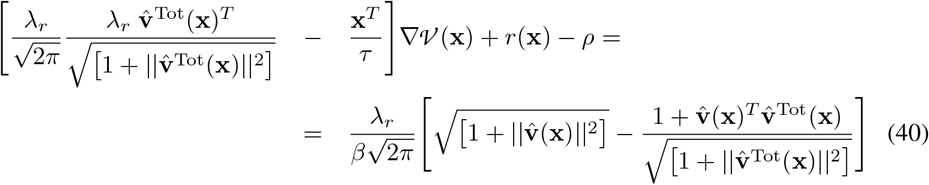

with boundary condition *𝒱* (**x**_*T*_) = *R*_*T*_ (**x**_*T*_).

For an example comparing the exponential and the sigmoidal non-linearities, see Figure Supp.1 (albeit for different types of terminal regions to also illustrate different regimes).

### A.5 Numerical solution of the Bellman Equation

To solve the Bellman Equation (BEq), we utilized the approach and neural network architecture proposed in [43] with modifications to suit our specific problem set. Unlike the approach in [43], where the boundary conditions are explicitly fitted, we adopt the methodology proposed in [42]. This approach involves modifications to the network architecture that inherently enforce the satisfaction of the boundary conditions. The general idea goes as follows. Let *𝒱* ^*^(**x**) be the value of the optimal policy, *S*_*θ*_(**x**) be a neural network with parameters *θ*, and Ω define a region in ℝ^K^, with **x**_*T*_ the terminal region. To ensure that the solution of the network satisfies the value on the terminal state, we can define our value approximation as 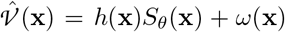, in which *h*(**x**_*T*_) = 0 but ∇ *h*(**x**_*T*_) ≠ 0, and *ω*(**x**) is a smooth extension of the function *𝒱* (**x**_*T*_), to Ω. The conditions on *h*(**x**) guarantee that the function *h*(**x**)*S*_*θ*_(**x**) is zero on **x**_*T*_ without imposing any artificial constrains on the derivative of 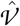 on the boundary. The condition on *ω*(*x*_*T*_) enssures that 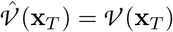.

For a broad array of problems, an architecture featuring a tanh activation function, three layers, and 250 neurons per layer effectively minimized the BEq. The network weights were initialized with Xavier’s initialization, or forcing the output of the network to be close to zero by scaling *S*_*θ*_(**x**) by a small value. The parameters were updated using the ADAM[58] optimizer with a learning rate of 10^−5^, adjustable based on the specific partial differential equation to address the expected smoothness of the Value function. The loss function was defined as the mean squared error of the differential operator in the BEq across a dynamically uniformly sampled set of points in Ω/{**x**_*T*_}. Notably, some BEqs, such as those with an exponential non-linearity, may have multiple solutions (e.g., both positive and negative). In our case, where only costs are considered, the only solution that has physical meaning requires that the maximum value occurs at the boundary. To enforce this, we integrated a regularizer into our loss function using the ReLU function to penalize non-boundary maxima, enhancing the accuracy and stability of the solution.

The batch of sampled points in which the loss is evaluated is drawn from a uniform distribution in Ω/ {**x**_*T*_} at every epoch. For most problems, we used a starting batch size of 2^11^, that increased linearly at a rate of 5 per 500 epochs throughout the training of the network. Typically, the network achieved satisfactory performance under 500000 epochs.

We observed that fitting the solution to the BEq can be challenging at low values of *β*. This difficulty arises because the control strategy must rely more heavily on the default dynamics of the spiking neural network to influence its activity. To address this issue, we developed a solution that involves initially solving the BEq at a high *β* value and then subsequently reducing *β* through a process of annealing, until it reaches the desired level. We observed that most annealing functions that decreases *β* sufficiently slowly will work. In particular we have used exponential decays so that the stepsize is proportional to the current value of *β*.

To train the networks, we utilized an NVIDIA RTX 4090 GPU. The training duration and the number of epochs necessary are highly variable, depending on the terminal state’s region and control problem parameters such as the cost of time, the cost of control, and the value at the terminal region. Consequently, network convergence times varied significantly; for some configurations, it took only 10 minutes to solve the BEq (figure Supp.1 **d**), while more complex problems required up to 5 hours (figure Supp.1 **b**). The primary factors influencing these variations are the number of epochs required to train the network and the floating-point precision used (32-bit or 64-bit). In certain settings, the control problem’s PDE solution is complex or features gradient discontinuities, necessitating higher floating-point precision to achieve an accurate approximation with a continuous network.

#### Algorithm 1

DGM with imposed Boundary Conditions

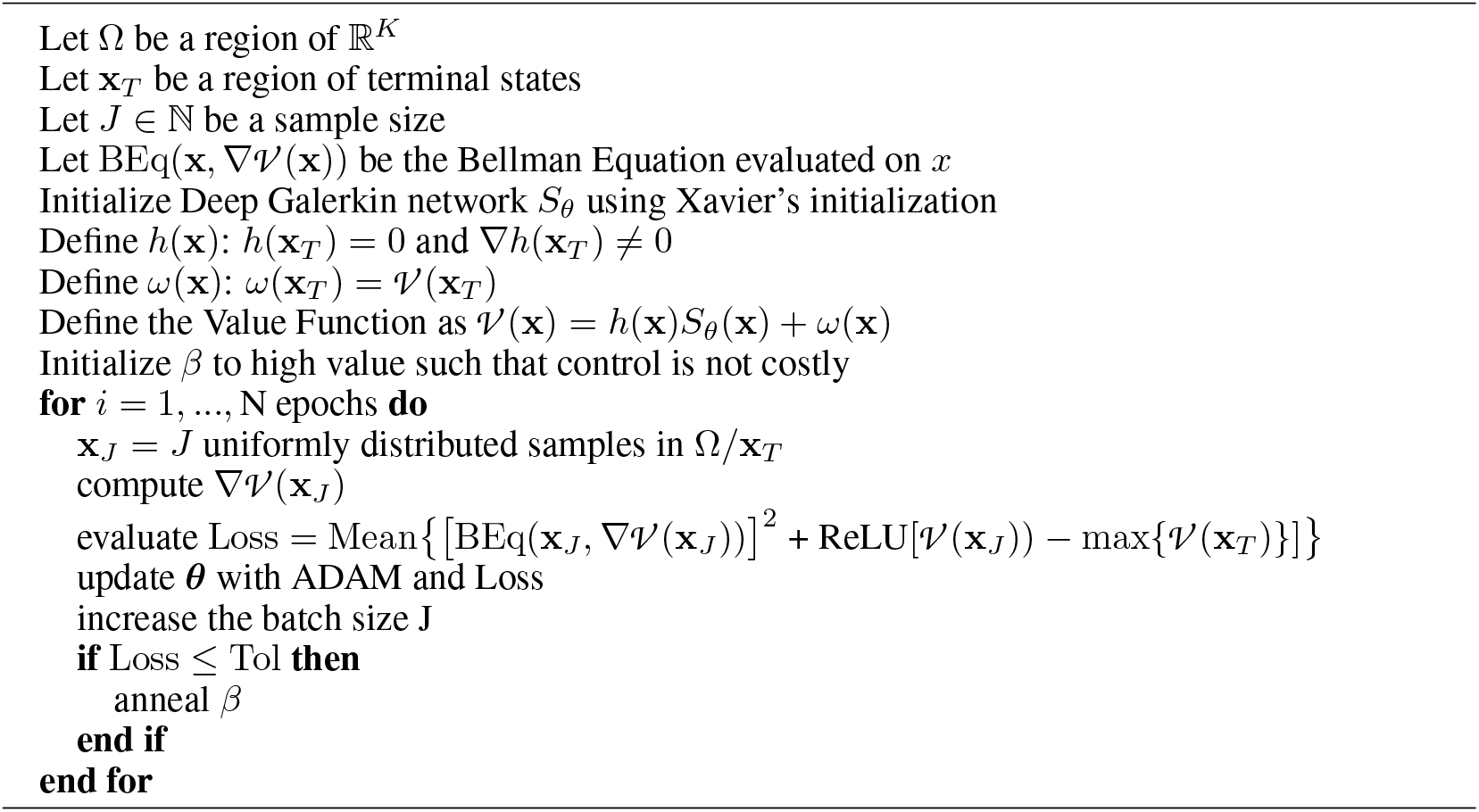

### A.6 On-policy-iteration method

In the main text, it was shown that the overlaps follow the stochastic dynamics (6):

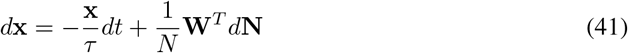

It can be shown that, in the limit of *N* growing to infinity, these dynamics can be approximated as an Itô process [24]:

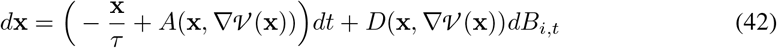

with *A*(**x**, ∇*𝒱* (**x**)) being the same quantity that was defined previously, and

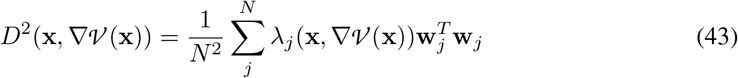

so *D*^2^(**x**, ∇ *𝒱* (**x**)) is of order 1*/N* and can be discarded. Thus, in the asymptotic regime, we are left with the following deterministic dynamics for the overlaps

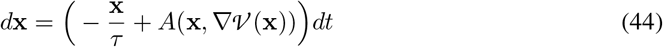

Based on this, we propose a new method for obtaining the optimal policy. The core concept involves proposing a policy, calculating the dynamics and accumulated costs under that policy, and using these costs to update the value function. Given that the control input to the network is *β*∇ *V* (**x**), a new policy can be derived for evaluation. This process is iterated until the value function and the policy stabilize, indicating convergence to the optimal policy. As in the main text, we have *B(* **x**, ∇ *𝒱* (**x**)) + *ρ* be the instantaneous cost.

We begin by defining an initial guess for the control, designed to ensure that all trajectories converge to the terminal states. Let *i* count iterations, and let us call *β* ∇ *𝒱*_*i*− 1_(**x**) the estimate for the policy based on the previous iteration, which is equal to the initial guess for the first iteration. We then uniformly sample points **x**_0,*j*_ in Ω/ {**x**_*T*_} (*j* denotes the index of trajectories). For each of the 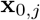 we compute the corresponding trajectory

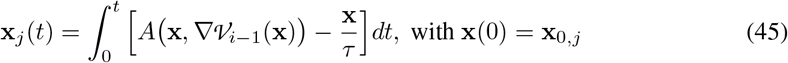

until **x**_*j*_(*t*) reaches **x**_*T*_. This sampling and computation process is repeated until the trajectories sufficiently cover Ω/ {**x**_*T*_}. For each trajectory, we compute the new estimate of the value by taking the path integral

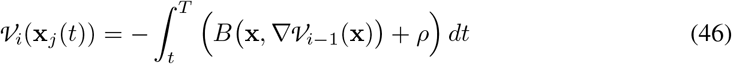

along the trajectory **x**_*j*_(*t*).

Having covered enough Ω/{**x**_*T*_ }for an accurate reconstruction, we interpolate this new estimate of value function using a cubic smoothing spline, which is advantageous because it allows for value interpolation anywhere within Ω and provides gradients that are smooth and inexpensive to evaluate. By definition, the control input associated with the new estimate of the value function is *β*∇*V*_*i*_. Using this updated estimate for the policy, we initiate the sampling process anew. We iterate the process until convergence is achieved, which is measured by the change in either the value or the gradients falling below a specified tolerance level.

We tested the algorithm with an Intel(R) Core(TM) i5-9600KF CPU (3.70GHz). Typically, it took between 10 and 20 iterations to reach convergence, as measured by a relative tolerance on the change of the value function and/or policy of less than 10^−4^. The time per iteration was quite variable, as we solved Equations 45 and 46 using “solve_ivp” from the scipy.integrate module (BFD method), which is an adaptive ODE solver that chooses the time step automatically. Therefore, depending on the stiffness of the problem, the trajectories might take longer to integrate, and also the number of trajectories per iteration might require adjustments in order to cover more densely problematic (i.e. steep gradient) regions. Nonetheless, the time per iteration was normally around 1-3 min, meaning that usually convergence was achieved in less than an hour. Typically, we use 500 or 1000 trajectories per iteration.

For an example illustrating both the Policy Iteration and the Deep Galerkin methods, see Figure Supp.2.

#### Algorithm 2

Policy Iteration

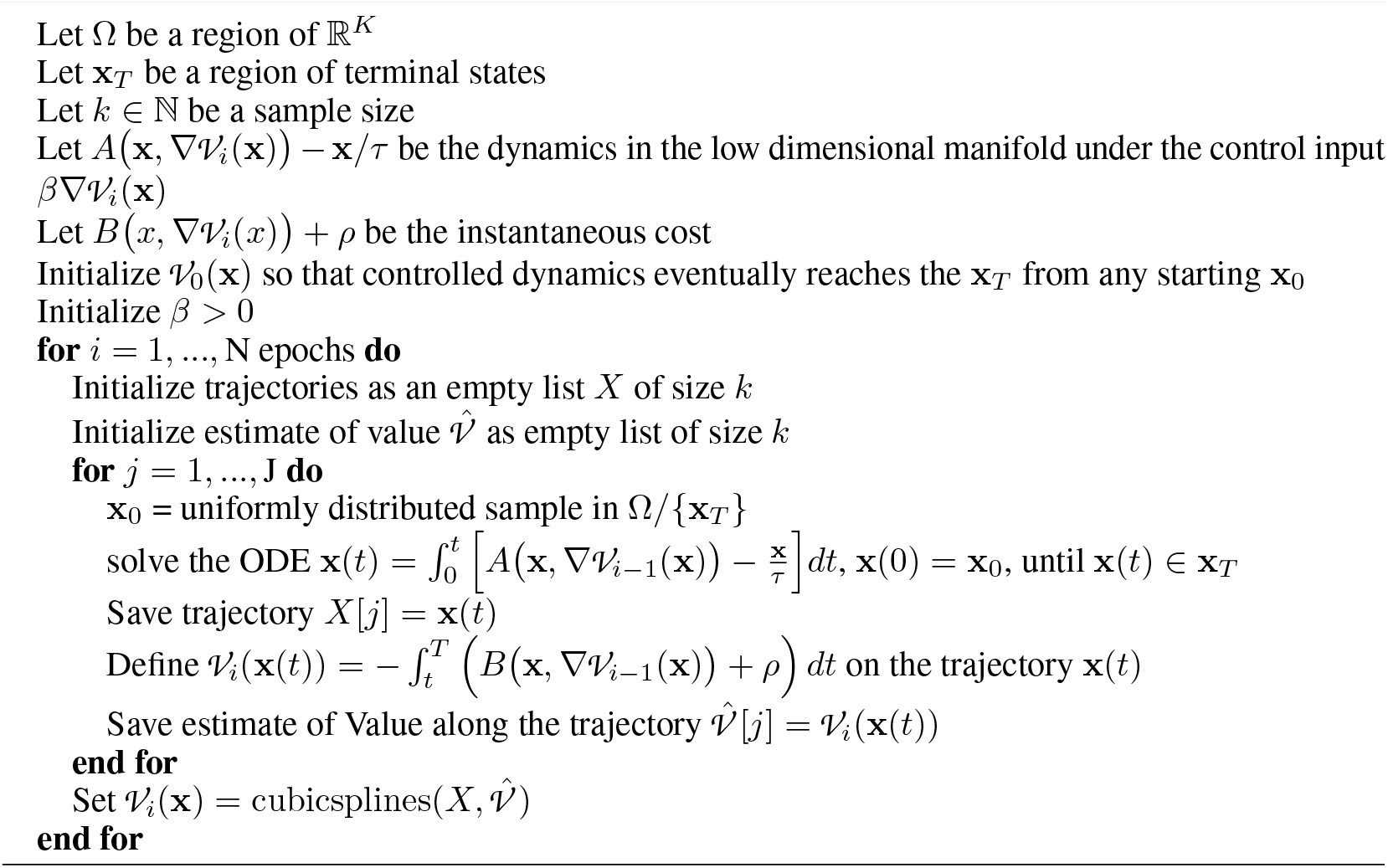

### A.7 Rank 1 network and values of the parameters used in the Figures

For the rank 1 network, 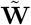 is a single vector 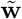 and **U** is a single vector 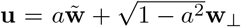 where *a* is the cosine of the angle between 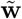 and **u**. Also, Σ is a scalar *σ*. In this case, Equations (3) and (4) read

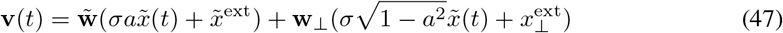

And

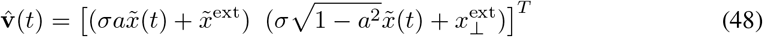

The values of the parameters used to generate all the figures are

*where 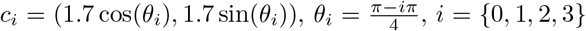, **x**^ext^ is a vector of components 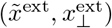 and **f** is a vector of components 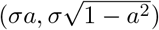. For all figures, the value at the terminal region **x**_T_= 0.

**Table 1:** Parameter values for the figures.

|  | params default dynamics |  |  |  | Params control |  |  |
| --- | --- | --- | --- | --- | --- | --- | --- |
| | $\lambda_r$ | $\tau$ | $\mathbf{x}^{\text{ext}}$ | $\mathbf{f}$ | $\beta$ | $\rho$ | terminal region |
| fig. 2 a, c | 2 | $\sqrt{2\pi}$ | (0, 0) | (0.5, 0.5) | NA | NA | NA |
| fig. 2 b, d, e, g | 2 | $\sqrt{2\pi}$ | (0, 0) | (0.5, 0.5) | 0.5 | 1 | $\{\mathbf{x} : (x_1 - 0)^2 + (x_2 - 1.7)^2 = 0.1^2\}$ |
| fig. 2 f, h | 2 | $\sqrt{2\pi}$ | (0, 0) | (0.5, 0.5) | 0.01 | 1 | $\{\mathbf{x} : (x_1 - 0)^2 + (x_2 - 1.7)^2 = 0.1^2\}$ |
| fig. 3 a | 2 | $\sqrt{2\pi}$ | (0, 0) | (0, 0.05) | 0.01 | 1 | $\{\mathbf{x} : (x_1 - 0)^2 + (x_2 - 1.7)^2 = 0.1^2\}$ |
| fig. 3 b | 2 | $\sqrt{2\pi}$ | (0, 0) | (0.5, 0.5) | 0.01 | 1 | $\{\mathbf{x} : (x_1 - c_{i,1})^2 + (x_2 - c_{i,2})^2 = 0.1^2\}^*$ |
| fig. sup. 1 b | 2 | $\sqrt{2\pi}$ | (0, 0) | (0.75, 0.5) | 0.01 | 1 | $\{\mathbf{x} : (x_1 - 1.25)^2 + (x_2 - 0.85)^2 = 0.1^2\}$ |
| fig. sup. 1 d | 1 | 1 | (0.7, 1.2) | (-0.05, -0.5) | 1 | 0.1 | $\{\mathbf{x} : x_2 = -4\}$ |

---

**Figure Supp.1:**
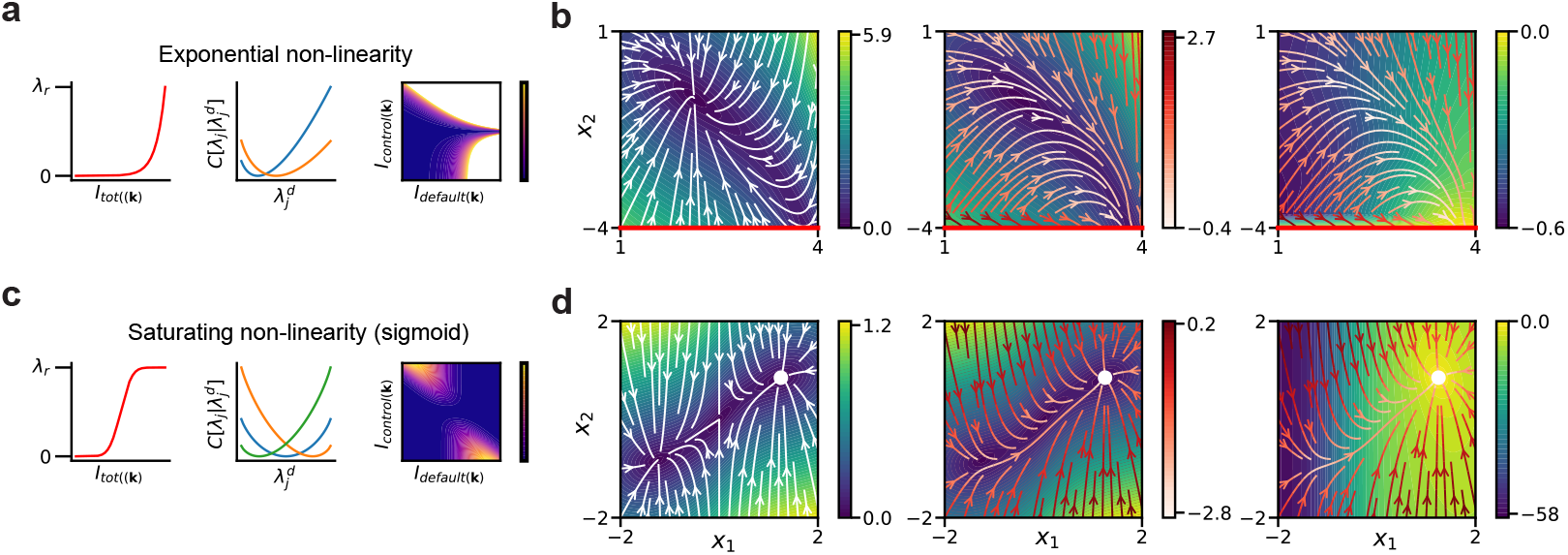
Comparison between exponential and sigmoidal firing rate non-linearities. **(a)** Firing rate non linearity (left), associated divergence as a function of the rates (middle) and as a function of the inputs (right). **(b)** Default dynamics (left), controlled dynamics with magnitude of the default in the background (middle) and with value in the background (right). The target region is the line *x*_2_ = − 4. **(c**,**d)** Same as **a**,**b**, but for a sigmoid non-linearity. In this case, the target is at one of the stable fixed points of the default dynamics.

**Figure Supp.2:**
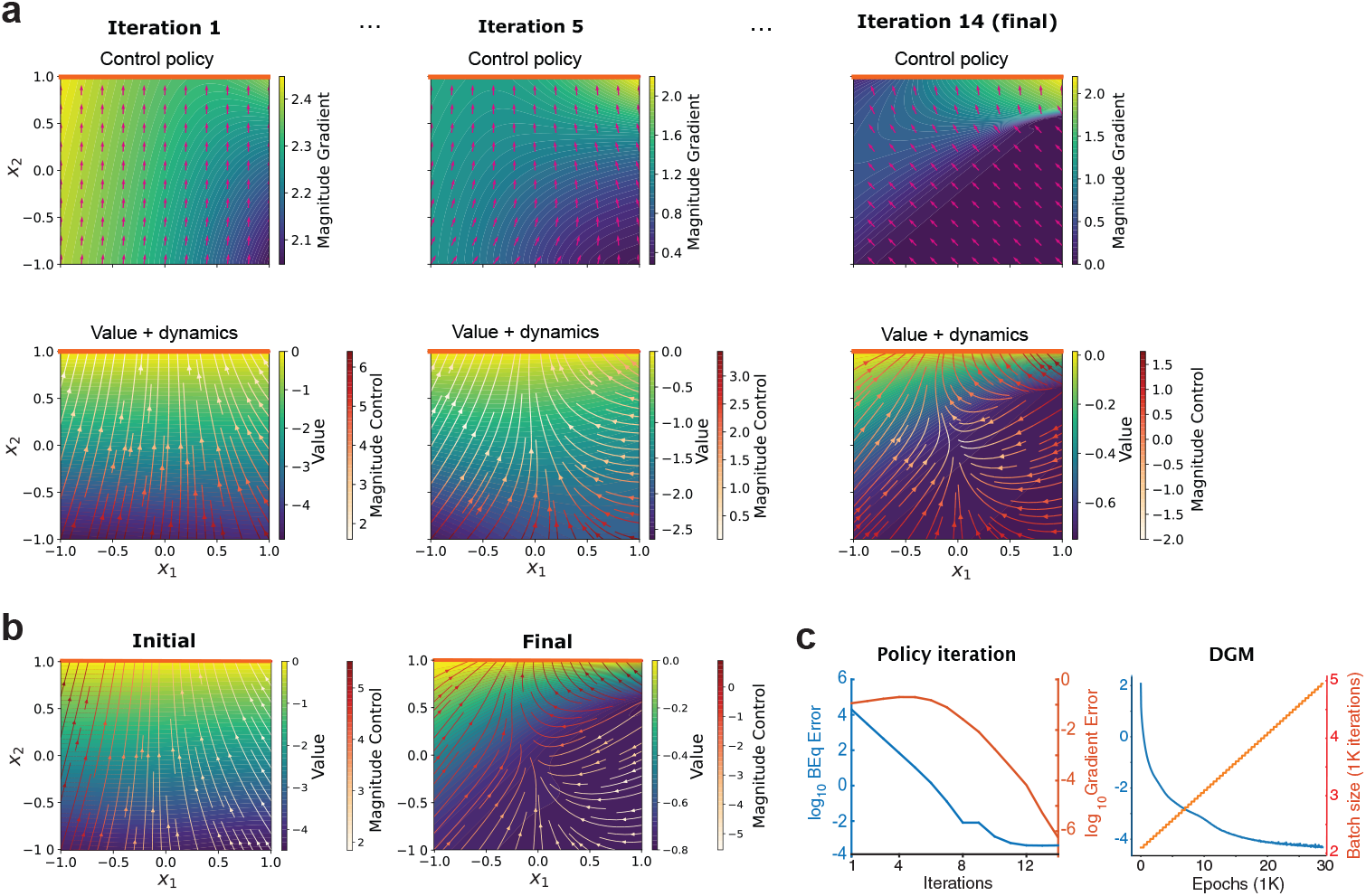
On-policy-iteration method and Deep Galerkin Method. **(a)** Example iterations for the on-policy-iteration method, for a target region located at the line *x*_2_ = 1. The first row shows the value gradient direction (arrows) and its magnitude (color), while the second row shows the streamlines for the controlled dynamics, with the Value on the background. **(b)** First and final states for the DGM, showing the corresponding controlled dynamics, with its magnitude represented in the color of the streamlines, and the value in the background. **(c)** On the left, error as a function of iteration number for the policy iteration method, measuring both the BEq error and the gradient error (mean absolute difference between iterations). On the right, BEq error and batch size as a function of epoch number for the DGM.

## Footnotes

1 We use [**M M**′] and [**M**; **M**′] to denote horizontal and vertical concatenation of matrices.

2 Because the spiking probability of different neurons is independent conditional on their membrane potential, it is the case that the control cost between two network-wide policies is equal to the sum between the control costs of each neuron in the network, so that 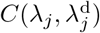 can be defined *per neuron*.

3 Note that, given the costs in Eq. (8), the value function has no explicit time dependence.

4 In principle, this maximization would require functional derivatives, since *𝒱* is a functional of λ. However, *𝒱* is already the optimal value, so it’s functional derivative with respect to λ is zero. The other terms in the equation depend only on λ itself, so the functional derivative can be evaluated as if it was a standard partial derivative.

